# Intracellular mechanisms of fungal space searching in microenvironments

**DOI:** 10.1101/391797

**Authors:** Marie Held, Ondrej Kaspar, Clive Edwards, Dan V. Nicolau

## Abstract

The underlying intracellular mechanisms involved in the fungal growth received considerable attention, but the experimental and theoretical work did not take into account the modulation of these processes by constraining microenvironments similar to many natural fungal habitats. To fill this gap in the scientific knowledge, we used time-lapse live-cell imaging of *Neurospora crassa* growth in custom-built confining microfluidics environments. We show that the position and dynamics of the Spitzenkörper-microtubules system in constraining environments differs markedly from that associated with unconstrained growth. First, when hyphae encounter an obstacle at shallow angles, the Spitzenkörper moves from its central position in the apical dome off-axis towards a contact with the obstacle, thus functioning as a compass preserving the ‘directional memory’ of the initial growth. The trajectory of Spitzenkörper is also followed by microtubules, resulting in a ‘cutting corners’ pattern of the cytoskeleton in constrained geometries. Second, when an obstacle blocks a hypha at near-normal incidence, the Spitzenkörper-microtubule system temporarily disintegrates, followed by the formation of two equivalent systems in the proto-hyphae – the basis of obstacle-induced branching. Third, a hypha, passing a lateral opening along a wall, continues to grow largely unperturbed while a lateral proto-hypha gradually branches into the opening, which starts forming its own Spitzenkörper-microtubule system. These observations suggest that the Spitzenkörper-microtubules system conserves the directional memory of the hyphae when they navigate around obstacles, but in the absence of the Spitzenkörper-microtubule system during constrainment-induced apical splitting and lateral branching, the probable driving force of obstacle-induced branching is the isotropic turgor pressure.

## Introduction

Filamentous fungi dwell in many and various geometrically-heterogeneous habitats, such as animal, human or plant tissue,^1,2^ decaying wood, soil, and leaf litter.^3,4^ The ecological ubiquity of filamentous fungi is, to a large extent, the consequence of their efficient search for space available for growth, often in mechanically-constraining geometries. Furthermore, because hyphae can grow for relatively long distances (millimetres) through media without, or with low level of nutrients, fungal space searching strategies need to be efficient even in the absence of chemotactic cues.

Extensive studies described several fundamental behavioural traits of fungal growth, including the directional growth of hyphae,^5-9^ regular branching,^10-12^ and negative autotropism^13,14^. These studies have overwhelmingly been performed on flat surfaces, usually made of agar, although the natural habitats of filamentous fungi typically present three-dimensional, constraining geometries. Opportunely, microfluidics devices, which have been interfaced with individual bacteria^15,16^ mammalian,^17,18^ and plant cells,^19,20^ and recently fungi,^21,22^ can be designed to mimic micron-sized, naturally-constraining habitats. Furthermore, the material of choice for the fabrication of microfluidics devices, Poly(dimethyldisiloxane), PDMS,^23^ is transparent, allowing the visualisation by microscopy techniques,^18,24^ and it is also permeable for O_2_, thus allowing *in vitro* studies in more realistic conditions.

Capitalising on the use of microfluidics technology, our previous studies,^25-27^ demonstrated, first for *Pycnoporus cinnabarinus*,^25^ and later for *Neurospora crassa,^26^* the very different behavioural traits of fungal growth in constraining geometries, e.g., up to ten times lower apical extension rates and distances between branches, compared with those on flat surfaces. The translation of the fungal space searching process into a mathematical formalism^25,28^ revealed that this strategy is analogous to a ‘master program’ with two ‘slave subroutines’: *directional memory*, whereby individual hyphae return to their initial growth direction after passing an obstacle forcing them to deviate from their course; and *obstacle-induced branching*, whereby branching occurs univoquelly when the hyphae encounter a solid obstacle blocking their growth. ‘Running’ this natural program resulted in a significantly more exhaustive exploration of the available space for growth than its alternatives,^25,26^ i.e., turning off either directional memory, or obstacle-induced branching, or both ‘subroutines’. It was also shown that the fungal space searching program can find exits in confining mazes quicker than some artificial space searching mathematical algorithms.^29^ However, given their behavioural focus, these studies could not offer explanations regarding the underlying ‘hard wired’ intracellular mechanisms responsible for the fungal efficient strategy for space searching in constraining geometries.

Studies regarding the intracellular mechanisms responsible for fungal growth, which used advanced fluorescence microscopy and were performed exclusively on non-constraining surfaces, revealed several essential processes.^7,30,31^ First, the positioning of the Spitzenkörper (a dynamic organelle complex) at the hyphal apex correlates with the direction of apical growth and the overall cell polarisation.^32-37^ Second, cytoskeleton dynamics (involving microtubules, actin, and protein molecular motors) mediate the directional, long distance transport of secretory vesicles from the body of the fungus towards the hyphal apex, carrying material for building the hyphal cell wall. While microtubule dynamics have been extensively studied,^38-43^ the understanding of the role of actin filaments is less developed and more recent.^44-48^ Third, the dynamic process of construction of hyphal walls results in an increase in stiffness from the apical to the basal regions.^38,41,43,49-52^ Finally, the gradients of ion concentration along the hypha and between the hyphal cytoplasm and the outside environment produce considerable turgor pressure, which provides a distributed internal driving force for fungal growth that is manifested primarily at the hyphal tip, and which allows the penetration through soft obstacles.^53-56^

Although the understanding of the growth-relevant intracellular processes, in particular the roles of Spitzenkörper, microtubules, and turgor pressure, is advanced and comprehensive, the large differences between the behavioural traits of fungal growth in non-constraining versus constraining environments suggest that the present knowledge requires an important upgrade. To elucidate the constrainment-specific intracellular process in fungi, in particular their role in directional memory and obstacle-induced branching, the present work used time-lapse laser scanning confocal microscopy to image the dynamics of growth of *Neurospora crassa*, in particular the dynamics of fluorescently labelled Spitzenkörper and microtubules, in confining microfluidics networks. The resulting insight into confined fungal growth is potentially relevant to various environmental, industrial, and medical applications, including fungal pathogenicity in animals and plants.

## Results

### Fungal growth on flat agar surfaces and in closed, non-constraining PSMS geometries

The experiments were performed in closed PDMS microfluidics structures comprising separate chambers for testing the responses of the intracellular mechanisms to confinement and constraining conditions (a description of the microfluidics experimental structures is presented in Figure 1, in the Methods section, and in Supplementary Figure SI 01; representative images of fungal growth in confining/constraining geometries are presented in Supplementary Figure SI 02). Because the vast majority of studies report on fungal growth experiments performed on agar, the first step in our study was to establish that the ‘internal’ control in our experiments, i.e., closed, but non-constraining, 100 x 100 x 10μm PDMS-made chambers, provides comparable growth conditions with those on agar. The three-way comparison of fungal growth, i.e., on agar, in closed/non-constraining conditions, and exposed to various level of constrainment, respectively, demonstrated that the conditions associated with the ‘external’ control on agar, and ‘internal’ control in microfluidics chambers, are similar. Supplementary Information presents a detailed discussion.

**Figure 1.**
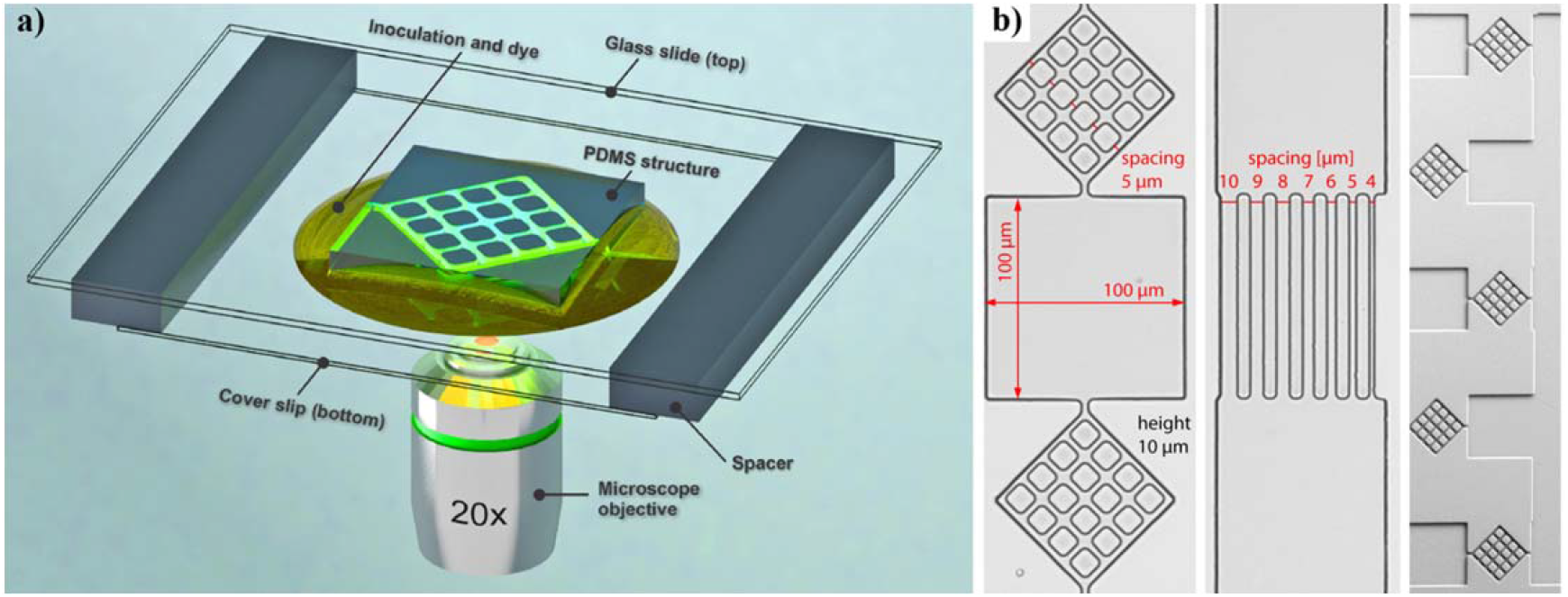
a) Schematic experimental setup for live-cell imaging of fungal growth in microfluidics structures on an inverted confocal microscope (not to scale). b) Micrograph images of the PDMS microfluidics structures constraining fungal growth. Left: chamber probing non-constraining growth (middle); diamond structure for probing lateral branching in constraining environments (top and bottom). Middle: channels for probing lateral branching following various constraining levels. Right: overall image of the entry to the chip, probing the response to collisions at shallow, and near-orthogonal angles, as well as corner responses.

Indeed, hyphal growth behaviour in PDMS non-constraining environments, including intracellular processes, is similar with that observed during ‘external’ control experiments on agar (Supplementary Figure SI 02). First, the cross-sectional apical profiles of *Neurospora crassa* hyphae are parabolic and symmetrical (Figure 2a for internal control; Supplementary Figure SI 04 for external control on agar). Second, the Spitzenkörper is centred at the hyphal apex (Supplementary Movie SI 01, and Supplementary Figure SI 05), with small periodic oscillations perpendicular to the growth direction (Supplementary Movie SI 02). Third, the longitudinal distribution of microtubule orientations is predominantly parallel to the longitudinal hyphal axis (Figure 2a for internal control; Supplementary Figures SI 04 and SI 06 for external control). For instance, in the apical regions, most microtubules (53%) deviate by less than 10° from the polarisation axis, and 84% deviate by less than 20°, with an overall mean deviation angle of 11.7° ± 9.5° (*n* = 453 microtubules measured in 20 hyphae, Figure 2b and Supplementary Movie SI 03). By contrast, in subapical compartments the angular deviations of microtubules are larger, i.e., 21% microtubules presenting a deviation of less than 10°, and 46% less than 20°, with an overall mean deviation angle of 26.8 ± 20.1° (*n* = 852 microtubules measured in 20 hyphae; Figure 2b and Supplementary Movie SI 04). Thus, the further away from the hyphal apex the microtubules are, the lower their alignment with the hyphal axis, as also reflected in the broadening of the distribution of microtubule deviations from the hyphal axis (Figure 2b for internal control; Supplementary Figure SI 04 and SI 06 for external control). A two-tailed test comparing the apical and subapical distributions of the alignment angles shows that the curves are non-identical (p < 0.0001).

**Figure 2.**
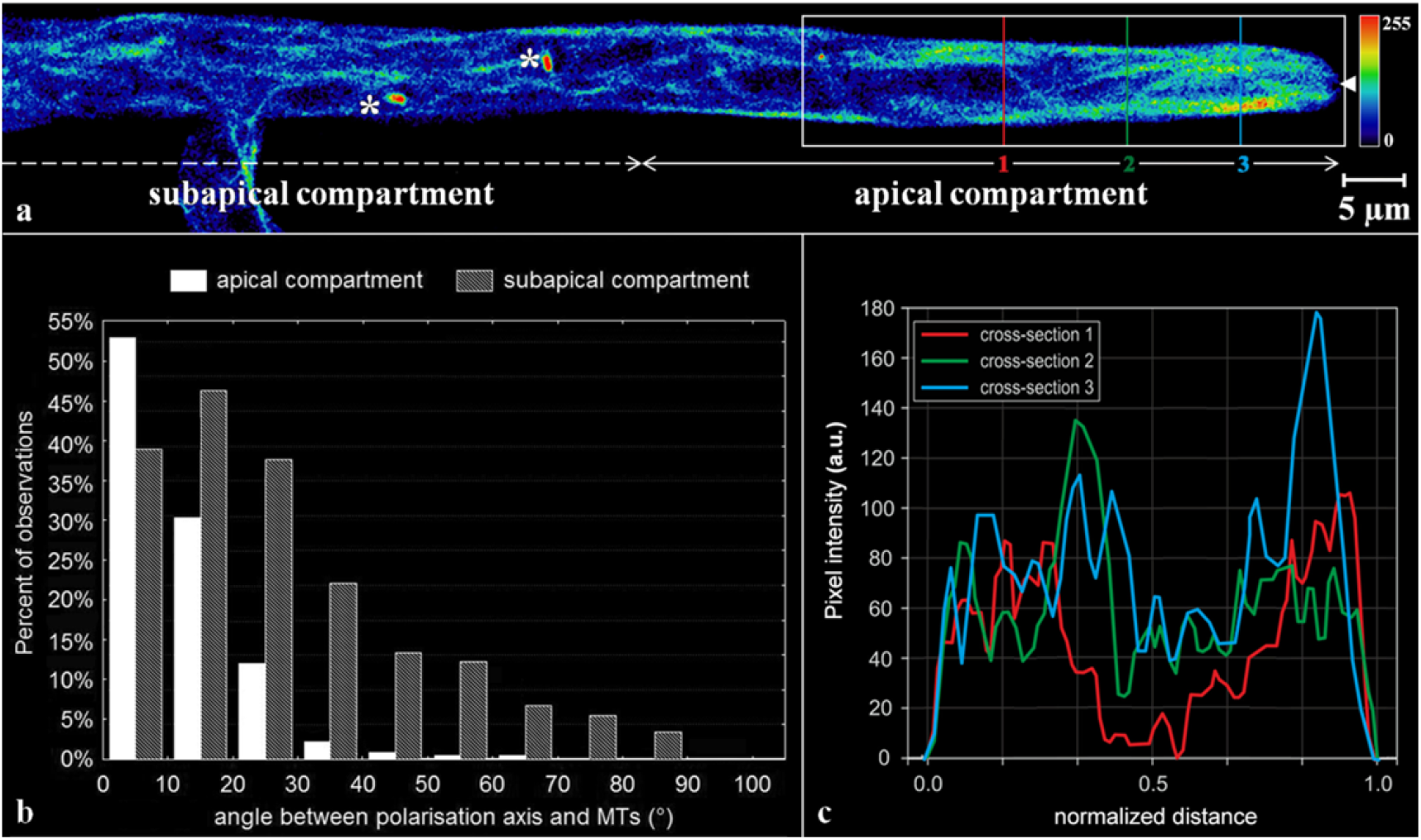
Spatial distribution of microtubules in *Neurospora crassa GFP* in non-confined environments. a) Single-plane fluorescence image of the GFP-tagged microtubules within a branched hypha. The microtubule alignment is predominantly longitudinal in the apical compartment (right) and less ordered in the subapical region (left). The colours represent the relative spatial density of microtubules (indicative colour map on the right). The asterisks indicate mitotic spindles, and the solid white arrowhead at the tip indicates the Spitzenkörper void. b) Histogram of microtubule deviations from the hyphal polarisation axis in the apical and subapical compartments (n = 852 microtubules measured in 20 hyphae). Longitudinal orientation is more pronounced in the apical region, where the microtubules also appear longer. c) Representative profiles of the microtubule density, calculated as fluorescence pixel intensities, along the vertical lines drawn across the hypha in a). The hyphal diameter, approximately 7 μm, was normalised to offset small variations at different sections through the apical compartment.

The lateral distribution of microtubules shows that while they populate the entire width of the fungal hypha, i.e., occupying both cortical and central cytoplasmic regions, the filament density is higher in the cortical region (Figure 2c for internal control; Supplementary Figure SI 07 for external control; Supplementary Table SI 02 and Supplementary Figure SI 08 present a statistical comparison between the two controls). The microtubules extend into the apical dome, displaying a characteristic microtubule-depleted zone in the distal central region that co-localises with the Spitzenkörper (Supplementary Movie SI 04). Long term imaging, e.g., between 5 to 10 min, showed that microtubules occasionally traverse the Spitzenkörper and frequently terminate at the apical cell wall. The estimated microtubule polymerisation rate is 26.4 ± 8.6 μm s^-1^ (*n* = 412 from 98 microtubules).

Finally, the lateral branching behaviour is also very similar on agar and in closed/non-constraining PDMS chambers, i.e., branching at approximately 45° with movement of microtubules into the daughter hypha (Supplementary Figures SI 09, SI 10 and Supplementary Movie SI 05), and central position and sizes of the Spitzenkörper (Supplementary Figures SI 10, SI 11 and SI 12). Additionally, experiments in non-constraining PDMS chambers enforced the growth of both parental and daughter branches on the same optical plane.

### Collision with obstacles at shallow angles

We then investigated how imposing geometrical constraints can affect hyphal growth. The geometry of the test structures (Figure 1b) presents a high density of various obstacles, such as corners, channels, entries and exits from large chambers, thus providing opportunities for *Neurospora crassa* hyphae to encounter various physical constrainments (also described before^25,26^). At shallow angles of approach, i.e., lower than 35° relative to the surface, the hyphae follows the contour of the obstacle, a process denominated as ‘nestling’. Nestling dynamics in *Neurospora crassa* hyphae (Figure 3, and Supplementary Movie SI 06) presents in three stages:

i. *Prior to nestling, i.e., prior to encounter with the wall*. As in non-constraining geometries, the hyphal profile is initially symmetrical and parabolic, with the Spitzenkörper centrally located at the hyphal apex, and the microtubules are symmetrically distributed. The consistency of the hyphal morphology prior to wall contact suggests the absence of any anticipatory, e.g., chemotaxis, sensing mechanism.
ii. *Nestling*. Hyphal morphology changes significantly when encountering a wall. First, hyphal growth follows the constrained path imposed by the obstacle, i.e., along the wall, in the direction of least deviation (Figure 3a, Supplementary Figure SI 13, top). Second, the longitudinal hyphal cross-section lose its parabolic symmetry, and becomes considerably skewed toward the wall. The hypha thus continues to progress in close contact with the wall, maintaining this skewed tip profile. Third, the Spitzenkörper markedly shifts away from its previously central apical location, towards the wall. This displacement persisted over longer distances, e.g., more than several hyphal diameters (Figure 3b, Supplementary Figure SI 13, bottom). Fourth, microtubules tend to gather near the inner edge of the hyphal bend (white arrow in Figure 3a) and towards the wall at the tip (Figure 3a, Supplementary Figure SI 14).
iii. *Return to non-constrained growth*. Upon reaching the end of the wall, the hyphae quickly recovers their original growth directions, within a distance approximately equivalent with the hyphal diameter. Additionally, the apex resume its symmetrical parabolic profile, and the Spitzenkörper simultaneously returns to a central position (Figure 3c; Supplementary Figure SI 15; Supplementary Movie SI 06 presents the complete time series) while the microtubules recover their symmetrical distribution.
iv. Within the spatial range of observation, spanning several chambers, each with a length of 100 μm, the accuracy in the recovery of the growth direction does not diminish over time, i.e., after successive bends through the device, or with the increase in distance from the initial branching point of the respective hypha (Supplementary Figure SI 16). Skewing of the apex during nestling is also constant over time.

**Figure 3.**
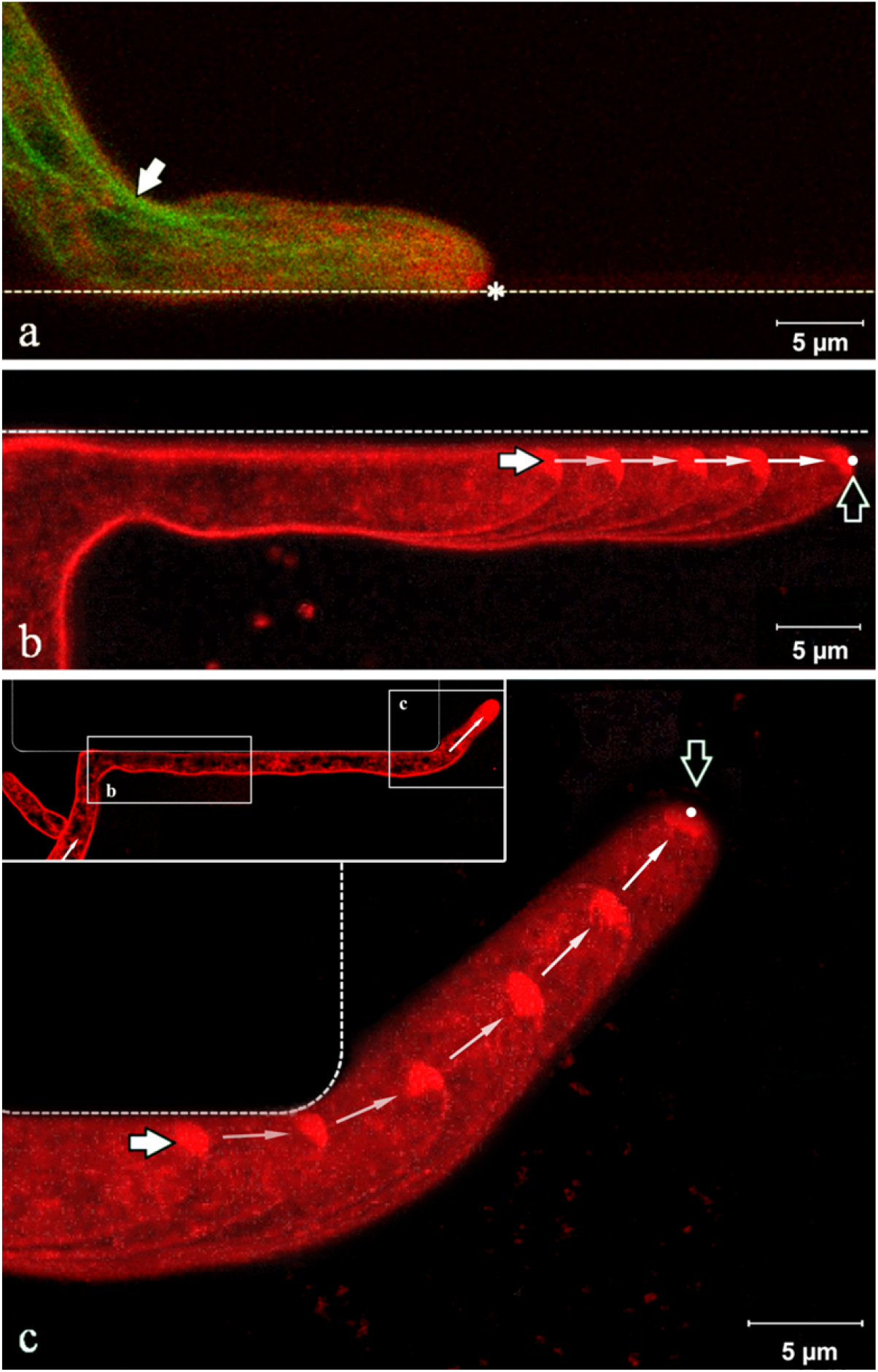
Fluorescence images of the Spitzenkörper (labelled with FM4-64, pseudo-coloured red) and microtubules (genetically tagged with GFP, pseudo-coloured green) in somatic *Neurospora crassa* hyphae nestling to a wall. a) Apical hyphal region growing along a PDMS wall (dashed line). The parabolic apex profile is skewed towards the wall. The Spitzenkörper (asterisk) is displaced from its usual central position at the apex as growth is obstructed by the wall. Microtubules follow the shortest path towards the Spitzenkörper (white arrow) and are displaced from the central median of the hypha. b) Trajectory of the Spitzenkörper, along the wall, displaced from the hyphal central axis, during nestling. The image overlays 5 snapshots taken over 4 min. The white and black arrows indicate the beginning, and the end, respectively, of the Spitzenkörper trajectory. c) Upon reaching the end of the wall, the hypha recovers its symmetrical parabolic profile and the Spitzenkörper gradually returns to the apex centre. The image represents an overlay of 6 images taken over 7.5 minutes, with the white, and the black arrows indicating the beginning and the end, respectively, of the Spitzenkörper trajectory. The images in b) and c) are from the same hypha at different times, as indicated in the inset of the c) image. The complete sequence of images can be viewed in Supplementary Movie SI 05.

The microtubule preferential distribution towards the wall opposing the direction of growth, which resulted in specific ‘cutting corners’ patterns, is also present when *Neurospora crassa* navigates channels that do not excessively constrain them, i.e., channel widths of 5 μm for a hypha diameter of 5-7 μm (Supplementary Movies SI 07, and SI 08).

### Branching induced by frontal collisions

Frontal encounters with a wall, defined as forming an angle greater than 35° relative to the obstacle surface, causes the splitting of the apices of *Neurospora crassa* hyphae (Supplementary Figure SI 17). Detailed and repeated imaging of this ‘hit & split’ process (n = 44 encounter events) put in evidence a process sequence with five stages (Figure 4, organised along stage number; also Supplementary Movie SI 09):

i. *Polarised approach, prior to encounter* (Stage 1, ‘Approach’ in Figure 4a1, b1, c1). The hypha approaches the wall, similar to the preliminary stage for nestling. The microtubules are orientated longitudinally, terminating at the apical cell region (Supplementary Figure SI 18, left panel).
ii. *From the moment of encounter until branching* (Stages 2-4, ‘Collision’ in Figure 4a2-4, b2-4, c2-4).

**Figure 4.**
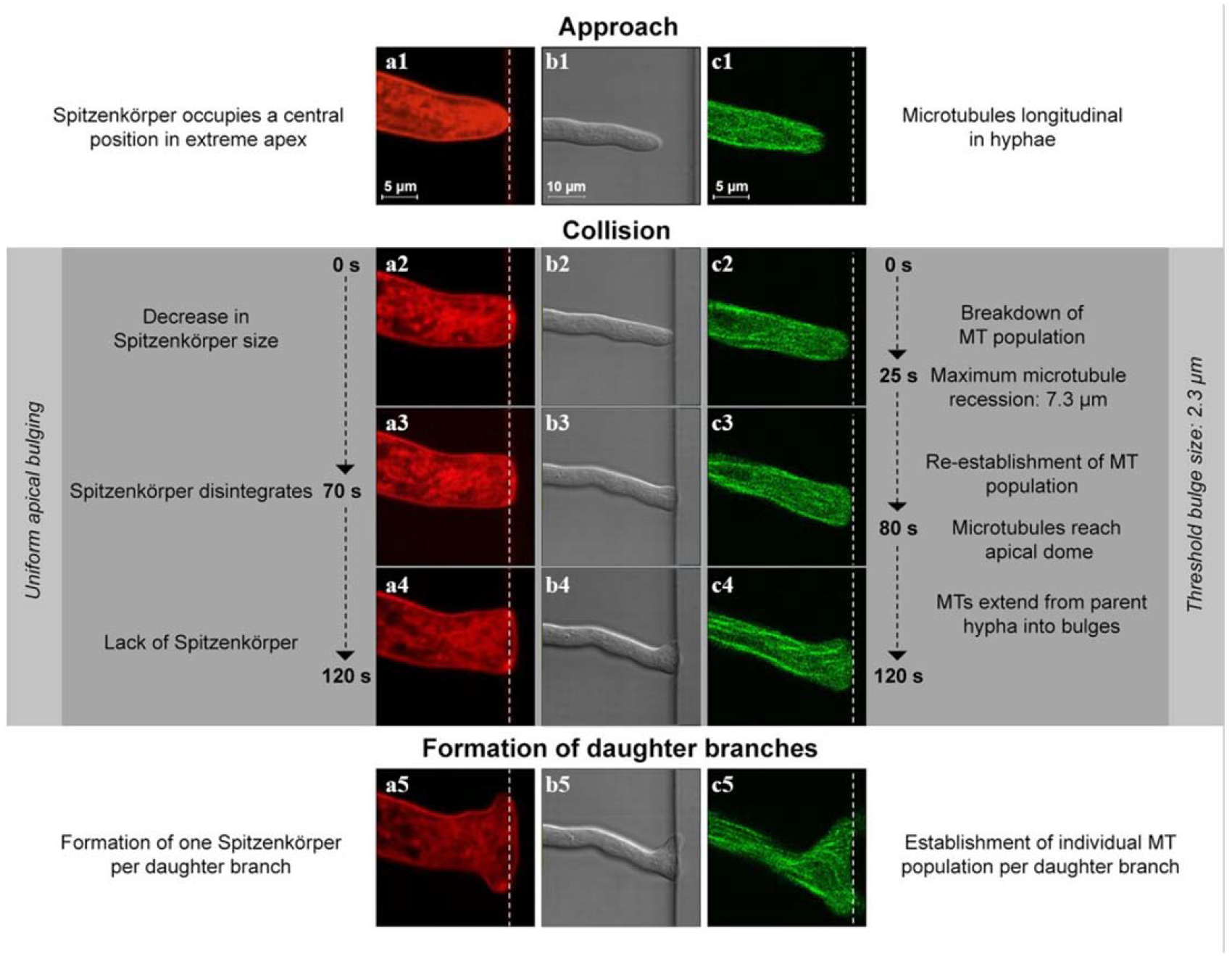
Stages during frontal obstacle-induced hit & split branching following collision with a PDMS wall (white dashed lines). Columns a and c represent the fluorescence images of the distributions of the Spitzenkörper (red) and microtubule (green), respectively, and column b represents the differential interference contrast image of a hypha. The hypha deforms the elastic PDMS slightly from its original position (b3, b4). During the approach (a1, a2), the Spitzenkörper is located at apex centre and the microtubules are organised longitudinally (c1, c2). Following the encounter, the Spitzenkörper shrinks (a2) and ultimately disappears (a3), and the microtubules temporarily recede from the apex region (c3, c4). Concomitantly, the apex grows uniformly (b3, b4). Finally, two new Spitzenkörper form in the daughter branches (a5) and the microtubules resume their extension towards both apices (c5).

In Stage 2 (Figure 4a2, b2, and c2), the obstacle blocks the extension of the apical cell in the growth direction. Apart from a slight wall deformation (due to PDMS elasticity), growth essentially continues orthogonally to the polarisation axis, resulting in lateral bulging in the apical region. Simultaneously, the microtubules depolymerise, with the filament ends receding rapidly from the apex (Figure 4c2, Supplementary Figure SI 18, second left panel). The average distance of filament end recession reached 7.3 ± 3.7 μm from the obstacle, at 25 ± 13 s after the collision. The Spitzenkörper does not retract longitudinally from the apical dome, but shrinks gradually (Figure 4a2).

In Stage 3 (Figure 4a3, b3, and c3), the hyphal profile continues to develop into two bulges. The total dissolution of the Spitzenkörper occurs towards the end of this phase, i.e., 70 ± 40 s after the initiation of the encounter (Figure 4a3). The microtubules resume their extension towards the apex, and 80 ± 36 s after collision, the microtubule population is fully recovered (Figure 4c3, Supplementary Figure SI 18, third left panel).

In Stage 4, just before branching is initiated, i.e., during the period when the hypha does not have a Spitzenkörper, the uniform apical extension continues laterally, following the constraining geometry, and the microtubules again extend to the extreme apical cell walls. The flexibility of the microtubules enabled their extension from the parent hypha into the nascent bulges, which ultimately resulted in extension perpendicular to the initial growth direction (Figure 4c4, Supplementary Figure SI 18, right panel).

*(ii) Branching*. In Stage 5, approximately two minutes after the orthogonal encounter with the wall, the uniform extension pattern changes to a bidirectional, polarised pattern, with the bulges reaching 2.3 ± 1.3 μm. The size of the buds immediately before forming new branches correlate moderately (r = 0.65, p < 0.05) with the initial diameter of the parental hypha. This change in polarisation pattern coincides with the nucleation of two smaller ‘daughter’ Spitzenkörper, one for each new growth direction (Figure 4a5). Additionally, independent microtubule populations develop within each branch to conclude the branching process (Figure 4c5).

Further evidence of the intracellular processes during the hit & split is provided in Supplementary Figure SI 19 for the Spitzenkörper trajectory; and in Supplementary Figure SI 20 for both the Spitzenkörper and the microtubules. Supplementary Figure SI 21 presents the history of the Spitzenkörper following the collision with an obstacle splitting the hypha in two branches.

Importantly, the disappearance of the Spitzenkörper also occurs when the hypha, pressed, then penetrated a PDMS wall (Supplementary Movie SI 10).

### Lateral branching from tightly constraining channels

*Neurospora crassa* branches with increasing frequency after prolonged bilateral constrain. Loose constrainment within channels with a width smaller than the hyphal diameter and without any side opening prevents branching over the entire channel length, but branching occurs almost immediately upon cessation of the confinement, e.g., at a channel opening (Supplementary Movie SI 11). However, if lateral openings are presented while a hypha passes through a narrow channel (Figure 5, Supplementary Figure SI 22, and Supplementary Movie SI 12), the intracellular processes responsible for directional memory and confinement-induced branching occur concomitantly.

**Figure 5.**
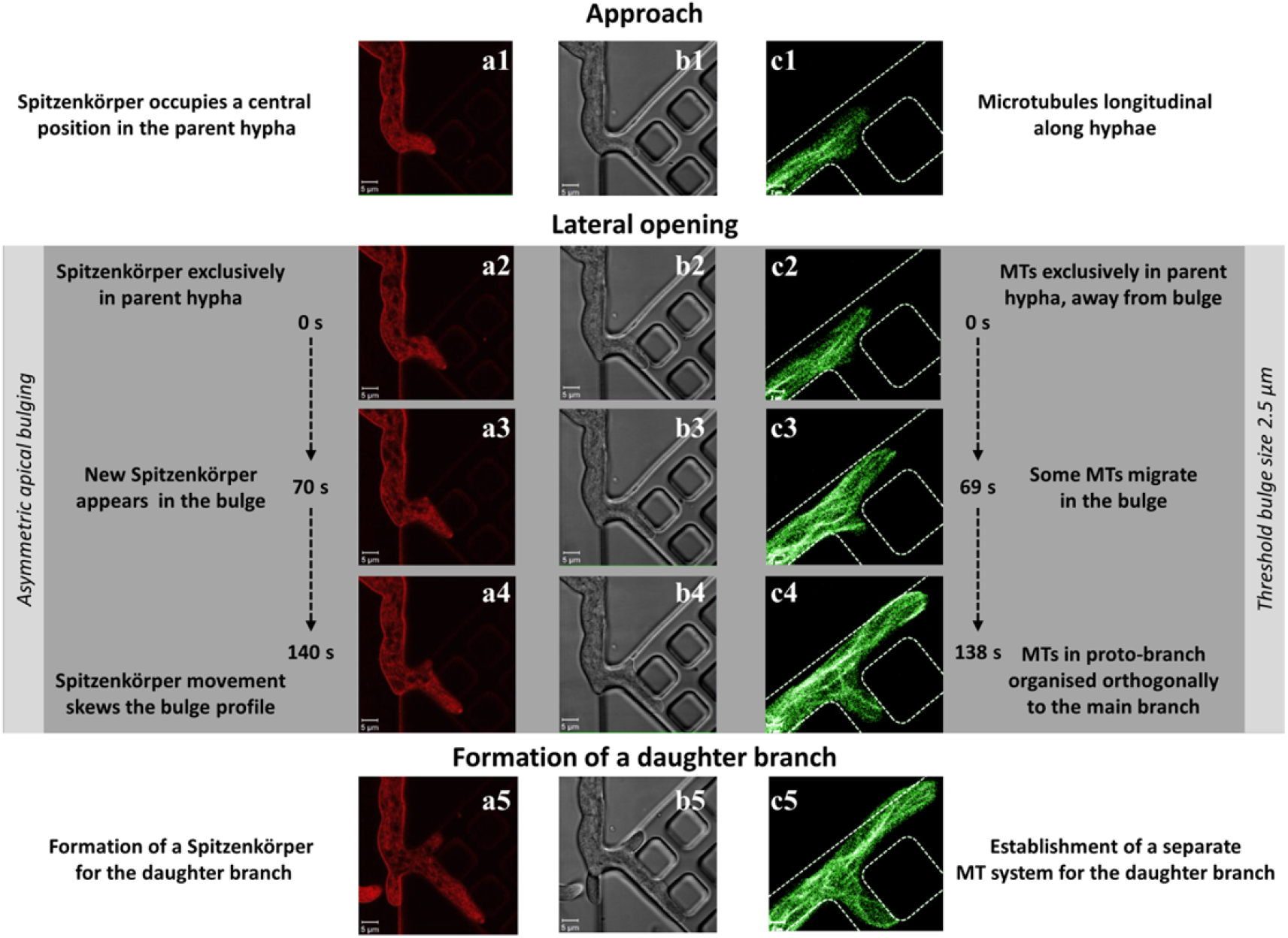
Stages of hyphal branching into a lateral channel (white dashed lines). Columns a and c represent the Spitzenkörper (red) and microtubule (green) distributions, respectively, and column b represents the differential interference contrast imaga of a hypha. The parent branch always preserves its Spitzenkörper. The Spitzenkörper is positioned closer to the wall, thus enabling directional memory. The formations of the Spitzenkörper and microtubule population in the daughter hypha are approximately simultaneous. While the cell wall senses the lateral gap, the formation of the daughter hyphae is delayed by the formation of the Spitzenkörper and microtubule population. The parent hypha in c) passes the intersection while the daughter branch forms orthogonally (c1-c2). Microtubules are initially distributed longitudinally in the parent hypha and do not extend into the bulge. Between frames (c3) and (c4), the microtubules start to extend from the parent hypha into the bulge, indicating the formation of the daughter hypha. The development of this branch is completed by the formation of an independent microtubule population (c5).

The growth and lateral branching proceeds in three stages (n = 20 hyphae):

i. *Entry and apical growth in the channel* (‘Approach’ in Figure 4a1, b1, c1). Directional memory manifests upon entering the confining channel (Figure 5a1 and b1), with the hypha extending along its initial growth direction, without turning into lateral channels. The microtubules are orientated longitudinally within the hypha (Figure 5c1).
ii. *Formation of a proto-branch* (‘Lateral opening’ in Figure 5a2-4, b2-4 and c2-4). When the apex encountered an intersection, the subapical hyphal regions gradually extend into the lateral opening. This orthogonal extension, aided by the plasticity of the cell wall, produces a bulge into the lateral space (Figure 5a2, b2, c2). Whereas the longitudinal microtubule orientation is initially preserved (showing no extension into the bulge, even after the hyphal apex had passed the lateral opening), at some point polarisation sets in (Figure 5c3), followed by an extension from the parent hypha into the developing branch (Figure 5c4). Approximately halfway through this process, this emerging branch forms its own Spitzenkörper (Supplementary Figure SI 23).
iii. *Development of a stand-alone orthogonal branch* (Figure 5a5, b5, c5). Subsequent development is characterised by the formation of an independent population of microtubules and an independent daughter hypha (Figure 5c5). Interestingly, features associated with directional memory appear early, e.g., the ‘cutting of corners’ pattern (Figure 5c5). This process occurs within only a few minutes from the initial crossing by the parental apex.

The directional memory in tightly-constraining environments manifests in the ‘cutting corners’ pattern for the microtubules in *Neurospora crassa* (Figure 6). Also, following the growth of a hypha towards a tight corner, the directional memory opposes a U-turn change in direction of growth, and instead, an orthogonal branch emerges near the apex of the parent hypha (Supplementary Movie SI 13).

**Figure 6.**
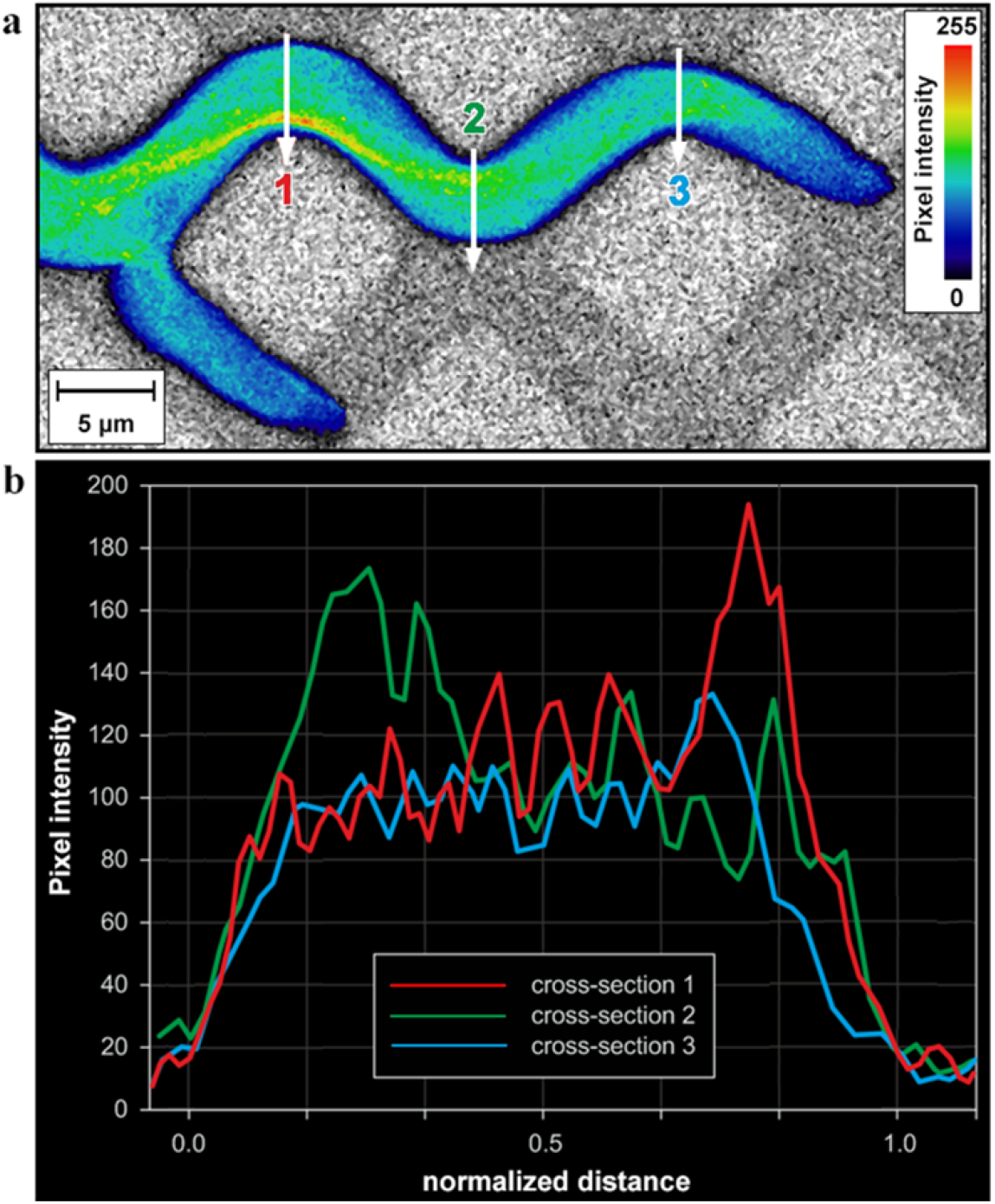
Spatial distribution of microtubules in *Neurospora crassa GFP* in confined environments. a) Single-plane fluorescence image of the GFP-tagged microtubules. The microtubule alignment largely follows the initial direction of growth at the entry in confining channels. The colours represent the relative spatial density of microtubules (indicative colour map on the right). b) Profile plot representing the microtubule densities, calculated as fluorescence pixel intensities, along the vertical lines drawn across the hyphal cross-section in a).

Additional evidence of the intracellular processes during the lateral branching in, or immediately after, tightly-constraining environments is provided in the Supplementary Information section: Supplementary Figure SI 22 presents the evolution of the microtubules during lateral branching in constraining environments, Supplementary Figure SI 23 presents the history of the Spitzenkörper in the same geometry; and Supplementary Figure SI 24 presents the evolution of the Spitzenkörper immediately after an exit from a tightly-constraining channel.

## Discussion

The intracellular mechanisms involved in fungal hyphal extension and branching have been comprehensively described in the literature, and they are still the subject of elaborate studies. However, the studies describing these intracellular processes have been so far invariably based on experiments performed on non-constraining artificial materials, almost always on flat agar surfaces. While this methodological choice is justified by the experimental difficulties associated with advanced microscopy, e.g., transparency of the substrate on which hyphal growth occurs, this experimental framework is dissimilar with many natural habitats of filamentous fungi, which comprise constraining geometries, and which are expected to interfere with the mechanisms of fungal growth.

Our previous studies, which used the visualization of the growth of filamentous fungi *Pycnoporus cinnabarinus*^25^, and later *Neurospora crassa^26,27^* in PDMS microfluidics structures, identified two behavioural traits, i.e., directional memory, and obstacle-induced branching, which distinguish the growth in confined spaces from that on flat surfaces, and which were proven to be efficient space-searching strategies.^25,28^

To this end, the present study aims to describe the differences, and the similarities between the intracellular mechanisms for hyphal extension and branching in non-constraining, and in geometrically constraining environments, respectively. The PDMS microfluidics structures used in this study enforce different levels of geometrical constraining of the hyphal growth: (i) virtually no mechanical constrainment, in closed, but non-constraining chambers; (ii) collision with a wall at shallow angles, leading to nestling; (iii) collision with a wall at near orthogonal angles, leading to ‘hit & split’ branching; and (iv) tight-lateral constrainment by narrow channels presenting either end-of-the-channel openings, or lateral ones.

### Intracellular mechanisms of fungal growth in non-constraining environments

We observed that the hyphal behaviour of *Neurospora crassa* in closed/non-constraining environments is similar to that observed during our growth experiments on agar, which correlate well with the reports in the literature. First, in our experiments the hyphae of *Neurospora crassa* present parabolic and symmetrical cross-sectional apical profiles (Figure 2a, Supplementary Figure SI 04), as also previously demonstrated and comprehensively described mathematically.^57-59^ Second, the central location of the Spitzenkörper at the hyphal apex (Supplementary Movie SI 01, Supplementary Figure SI 05) was described early in classical studies.^60^ Also, the observed oscillations orthogonal to the growth direction (Supplementary Movie SI 02) are consistent with previously reported results.^6^ Third, the general orientation of the microtubule parallel to the longitudinal hyphal axis (Figure 2a, Supplementary Figure SI 06, SI 07, SI 08, and Supplementary Table SI 02), and their accumulation towards the apical region, correlate well with those observed previously.^38,40,41^ Forth, the microtubule polymerisation rate (26.4 ± 8.6 μm s^-1^) is consistent with previously reported results obtained for hyphal growth on agar.^38^

To conclude, a high degree of similarity exists between the growth behaviour and related intracellular processes, for our experiments in closed/non-constraining chambers, and on agar, and for results reported in the literature. Consequently, the experiments in large microfluidics chambers are valid benchmark controls for the further assessment of the constrainment on fungal growth.

### Intracellular mechanisms responsible for directional memory during nestling

In general, the extension of hyphae over a flat surface follows a direction determined at the initial branching point, usually at an angle of approximately 45° from the parent hypha (Supplementary Figure SI 09). In contrast, in constraining geometries the hyphae are obligated to grow in the direction imposed by obstacles, or if space is available, to branch. We previously showed^25,26^ that once the obstacle was overtaken, the hyphae recovered their initial growth direction to within approximately 20°. This ‘directional memory’ persisted even over distances longer than ten times the hyphal diameter, regardless of the number of encountered collisions. Interestingly, the directional memory was demonstrated in both *Pycnoporus cinnabarinus*^25^ and *Neurospora crassa*,^26^ but not in the cytoskeleton-defective *Neurospora crassa* ro-1 mutant.^26^ This observation suggests that the dynamics of cytoskeleton play a central role in the maintenance of directional memory in constraining geometries.

Our results in non-constraining environments, i.e., on agar, or in 100 x 100 x 10 μm PDMS-made chambers, confirm earlier observations that hyphal growth follows the positions taken by the Spitzenkörper.^6^ However, while this observation remains valid when hyphae overtake obstacles during nestling, it also requires important qualifications. Indeed, when a hypha encounters a barrier at shallow angle of contact, thus overtaking the obstacle via sliding, the Spitzenkörper operates like a compass pointing in the direction the hypha had before the encounter (Figure 3, Supplementary Movie SI 05, and Supplementary Figure SI 13). One possible explanation for this, until now unreported, process is that the pressure applied to the hyphal wall due to the mechanical contact with the obstacle results in an internal signal, which triggers the consolidation of the hyphal wall in the zone of contact. This process would require the positioning of the Spitzenkörper off-axis and pressing on the contact point between the hyphal wall and the obstacle (as confirmed in additional experiments, e.g., Supplementary Figure SI 24). Furthermore, the off-axis position of the Spitzenkörper translates into a skewed architecture of the microtubule cytoskeleton, which present a characteristic pattern of ‘cutting corners’ (Figure 3a), in particular when the directional memory manifests in hyphae overtaking corners in a meandered channel (Supplementary Figures SI 25, SI 26, SI 27 and SI 28; and Supplementary Movie SI 14). This effect is even more remarkable considering that the microtubules must pass initially and/or eventually through narrow septa, which are centrally located on the median line of the hypha^61,62^ (Supplementary Figures SI 29, SI 30; and Supplementary Movie SI 15). The synergy between the compass-like function of the Spitzenkörper, subsequently enforced by the preferential positioning of the microtubules along a line approximating the initial direction of hyphal growth, appears to constitute the underlying intracellular mechanism for directional memory, which was observed for distances at least one magnitude longer than hyphal diameters (the hyphal trajectories in the Supplementary Movie SI 07 and Movie SI 08 are longer than 100 μm; and the distances in the Supplementary Figure SI 15 are several hundreds of μm).

More detailed experiments regarding the role of F-actin structures, i.e., actin rings, patches, and cables,^46^ which are more difficult to visualise than microtubules,^46,47^ would reveal their potential role in directional memory. However, because actin cables are co-localised near the Spitzenkörper and behind actin rings, it is expected that the role of actin is limited, at least in the long-range aspect of directional memory.

### Intracellular mechanisms responsible for obstacle-induced branching during hit & split

Our previous experiments with *Neurospora crassa^26^* showed that the constrainment effects affecting fungal growth in various microfluidics structures results in shortening the distance between hyphal branching points by a factor between 5 and 10 (the growth rate also decreases ten-fold). We also observed^26^ that the branching of *Neurospora crassa* when colliding with an obstacle at near-orthogonal angles occurred at the apex of the hypha, immediately following the contact between the hyphae and the constraining structure. This hit & split branching behaviour contrasts the one presented by *Pycnoporus cinnabarinus*^25^, which branches at a considerable distance behind the hyphal apex.

The observed Spitzenkörper dynamics in hit & split branching presents similarities with the processes during apical branching of *Neurospora crassa* on agar, which was studied in detail, using phase contrast video-enhanced light microscopy,^63^ and confocal fluorescence microscopy.^40^ For instance, both the disappearance of the parent Spitzenkörper, after a microtubule contraction from the apex region and the nucleation of the two daughter Spitzenkörper centres were also observed in the apical branching of *Neurospora crassa* on agar^63^. More specifically, in internally-triggered apical branching on agar^63^ the Spitzenkörper retracts 12 s after cytoplasmic contraction from the apex which precedes the branching, and disappears after another 47 s. Later, 45 seconds after the start of isotropic, uniform, albeit slower growth of the parental and daughter hyphae, one Spitzenkörper nucleates, followed by a second one approximately 7 seconds later, leading eventually to the establishment of two new branches. By comparison, in our observations of hit & split branching (Figure 4, Supplementary Figure SI 17, SI 18), the Spitzenkörper is missing for an average time of 50 seconds (n = 44). Moreover, the observed decrease in Spitzenkörper size, its subsequent disappearance, and the assembly of two new daughter Spitzenkörper centres away from the parent Spitzenkörper position, form a typical sequence of events that also occurs naturally in apically branching fungi, e.g., *Sclerotinia sclerotiorum^64^*

The dynamics of the Spitzenkörper in hit & split branching also presents significant differences when compared with that manifested during branching in non-constraining environments, both observed in our experiments on agar, and presented in the literature^63^. First, on homogeneous agar substrates the branching of *Neurospora crassa* hyphae occurs predominantly laterally, not apically.^63^ However, we observed that apical branching is the prevalent process in hit & split branching. Second, the apical extension stalls during the absence of a Spitzenkörper in *Sclerotinia sclerotiorum*,^64^ and is notably reduced in *Neurospora crassa* branching apically on agar.^63^ In contrast, this delay is not observed in our experiments with *Neurospora crassa* colliding near-orthogonally with a wall. We attribute this difference in the extension rates between during hit & split branching, and apical branching in non-constraining environments, respectively, to different trigger mechanisms. For example, an apical split can occur on agar few minutes after receiving an intracellular signal, whereas the immediate response of *Neurospora crassa* following a frontal collision with an obstacle, as observed in the present study, can be the result of a cascade of very localised, mechanical contact-induced processes.

The behaviour of the microtubules in apical and lateral branching on agar is similar,^40^ but it is markedly different during the hit & split response. In the apical or lateral branching on agar, the microtubule population is relatively constant throughout the branching process, whereas a hit & split response appears to trigger microtubule dissolution (Supplementary Figure SI 20, SI 31). No microtubules are observed to be associated with the cell wall, or to break by bending. The microtubules disintegrate, which created a gap of a few micrometres between the microtubule end points and the bulging cell wall. Furthermore, when a hypha encounters a corner (Supplementary Movie SI 10), branching occurs faster, producing only one branch on the side of the parent hypha, as allowed by the confining geometry. In this instance, the resulting budding branch does not display an initially discernible microtubule population, suggesting that the association of microtubules with the apical cell wall is not a prerequisite for selecting a branching site, as observed for the lateral branching in non-constraining environments,^40^ but which could be alternatively explained by cell-wall deformation driven by isotropic turgor pressure.

The present study also found that the extension rates of mature hyphae depends on the microtubule dynamics, as also reported before.^38^ However, in contrast to the ‘internal disruption’ of the microtubule dynamics by mutations,^38^ in the present study the dynamics is disrupted externally by enabling encounters with the constraining geometry. The filaments respond collectively via an asymmetrical distribution or population breakdown, thereby disrupting the supply of vesicles to the Spitzenkörper within the apex. This asymmetry is maintained over a long period during nestling (Supplementary Figure SI 32), but is short-lived for obstacle-induced breakdown dissolution (Supplementary Figure SI 31). The initiation of the recruitment sites for the morphological machinery in bilateral constrainment and frontal collision occurs in the absence of clear involvement of microtubules, and instead primarily through cell-wall deformation. By elimination, turgor pressure is the most likely cause of the branching that follows hyphal collision with obstacles at near-orthogonal angles.

Similarly to nestling, the role of actin in the hit & split branching remains to be established by further experimentation. However, as it was shown, for two yeast species,^65^ and for *Neurospora crassa,^66^* that actin is not present at the tip of invasive hyphae, i.e., those pressing against agar, i.e., in conditions similar to our experiments (Supplementary Movie SI 08 and SI 09). Consequently, it is expected that the contribution of actin is minimal to hit & split branching.

### Overlap of intracellular mechanisms of directional memory and obstacle-induced branching during lateral branching

*Neurospora crassa* predominantly branches laterally in non-constraining conditions, e.g., on flat agar surfaces, as observed in this study, and reported comprehensively in the literature, as well as in large PDMS chambers. Lateral branching also occurs in tightly-constraining microfluidics channels, but this process presents both similarities and differences when compared with the lateral branching on non-constraining conditions.

At the beginning of lateral branching in non-constraining geometries, cortical microtubules are associated with the cell wall at the location of the developing lateral branch. Upon further extension, the microtubules gather and bent considerably. The severed ends then migrate into the branch and resume polymerisation. These observations correlate well with those reported regarding the intracellular processes during lateral branching on flat agar surfaces.^40^ Importantly within the present context, in tightly-constraining channels, the original Spitzenkörper remains intact in the parental hypha during lateral branching, and a new Spitzenkörper is created independently within the ‘daughter branch’, which is also observed in lateral branching in non-constraining conditions.^63^

The most evident difference between the lateral branching in tightly-constrained geometries and that on flat surfaces is that the branching place and frequency is dictated by the availability of lateral space, rather than triggered by an internal clock, as it appears to be the case for non-constraining conditions. Moreover, there is a close temporal correlation between the presence of a constraining geometry and the lateral branching in a tightly-constraining channel. Also, the growth direction of the lateral branch is enforced by the axis of the available space for branching, e.g., orthogonal in Figure 6 (also Supplementary Figure SI 22, SI 23) rather than the usual approximately 45° in non-constraining conditions. These observations suggest a critical role of the isotropic turgor pressure in initiating the lateral branching events. Additional evidence suggesting the role of turgor pressure in lateral branching in our study is that *Neurospora crassa* branches commonly and almost immediately after an exit from a bottleneck (Supplementary Movie SI_11),^26^ but not *Pycnoporus cinnabarinus*,^25^ with a mechanically-strong hyphal wall needed for the penetration of wood litter.

Finally, our findings regarding branching in constraining environments differ from those of a previous study^38^ involving the same genetically tagged *Neurospora crassa* strain, but with lateral branching occurring on non-constraining flat agar surfaces. In our study, no cortical microtubules are observed bending or being shattered. Cell-wall deformation preceds microtubule extension from the parent hypha into the nascent bud, making it appear as the dominant element in the chain of events leading to branch formation. The bulging of the cell wall into an intersection of channels also precedes the formation of a daughter Spitzenkörper (Supporting Figure SI 23), suggesting that Spitzenkörper nucleation occurs after the initiation of branching.

The lateral branching in tightly-constrained channels presents the concomitant play of directional memory driving the growth of the parental hypha, which is modulated by the Spitzenkörper-microtubules system, and of obstacle-induced branching driving the growth of the daughter hypha, whose initiation appears to be the result of the turgor pressure overcoming the mechanical strength of the hyphal wall in locations where the back support of the constraining wall ceases due to a lateral opening.

### Intracellular mechanisms of directional memory and obstacle-induced branching

The use of time-lapse confocal fluorescence microscopy applied to the observation of growth of *Neurospora crassa* in various constraining microfluidics environments put in evidence substantial differences in the intracellular processes involved in the fungal search for available space for hyphal growth, compared with those manifested in non-constraining conditions. These differences are summarised in Table 1.

**Table 1.** Comparison of intracellular processes involved in the growth of *Neurospora crassa* in open and constraining environments. Present study*. CNC = “confined, but non-constraining”.

| Growth | Hypha | Spitzenkörper | Microtubules |
| --- | --- | --- | --- |
| **Non-constraining geometries [various references, \*]** |  |  |  |
| Single hypha | <u>Profile</u>: parabolic, laterally-symmetrical<br><u>Source</u>: agar<sup>57-59,*</sup> and CNC* | <u>Location and dynamics</u>: central, at the hyphal apex; permanently present<br><u>Source</u>: agar<sup>60,*</sup> and CNC* | <u>Orientation</u>: parallel to the hyphal axis<br><u>Distribution</u>: axially symmetrical<br><u>Dynamics</u>: population relatively constant<br><u>Source</u>: agar<sup>39-41,*</sup> and CNC* |
| Lateral branching | <u>Occurrence</u>: statistically-regular<br><u>Angle</u>: approx. 45°<br><u>Profiles</u>: parabolic for both parental and daughter hyphae<br><u>Apical extension</u>: reduced during branching<br><u>Source</u>: agar<sup>63</sup> and CNC* | <u>Location and dynamics</u>: central, at the hyphal apices; permanently present in parental hypha, early appearance in the daughter hypha<br><u>Source</u>: agar<sup>63</sup> and CNC* | <u>Orientation</u>: parallel to hyphal axes<br><u>Distribution</u>: axially symmetrical<br><u>Dynamics</u>: population relatively constant<br><u>Source</u>: agar<sup>63</sup> and CNC* |
| Apical branching | <u>Occurrence</u>: statistically-regular, but rare<br><u>Angle</u>: V-shaped, approx. 45°<br><u>Profiles</u>: initial round-up for the twin hyphae<br><u>Apical extension</u>: reduced during branching<br><u>Source</u>: agar<sup>63</sup> | <u>Location and dynamics</u>: retracts from the apex, disappears, then two Spitzenkörper centres emerge at the centres of hyphal apices<br><u>Source</u>: agar<sup>63</sup> | <u>Orientation</u>: parallel to hyphal axes<br><u>Distribution</u>: axially symmetrical<br><u>Dynamics</u>: population relatively constant<br><u>Source</u>: agar<sup>40,67</sup> |
| **Constraining geometries [\*]** |  |  |  |
| Nestling | <u>Occurrence</u>: triggered by contact at shallow angles<br><u>Angle</u>: change of direction as dictated by the wall<br><u>Profiles</u>: skewed off-axis, towards the wall<br><u>Apical extension</u>: unchanged | <u>Location and dynamics</u>: off-axis location, pressing against the obstacle, return to central position after passing the obstacle | <u>Orientation</u>: aligned off-axis,<br><u>Distribution</u>: axially asymmetrical, ‘cutting corners’<br><u>Dynamics</u>: population relatively constant |
| Hit & split | <u>Occurrence</u>: triggered by near-orthogonal collisions<br><u>Angle</u>: T-shaped branching, at ~180°<br><u>Profiles</u>: triangular, then progressively parabolic<br><u>Apical extension</u>: constant during splitting | <u>Location and dynamics</u>: disappears during splitting of parental hypha; then two Spitzenkörper centres form centrally at the apex of twin branches | <u>Orientation</u>: random close to the splitting<br><u>Distribution</u>: random close to the splitting<br><u>Dynamics</u>: substantial dissolution, then formation in twin hyphae |
| Branching in/after tightly-constraining channels | <u>Occurrence</u>: triggered by availability of free space for branching<br><u>Angle</u>: dictated by the geometry<br><u>Profiles</u>: parabolic for parental hyphal circular, then increasingly parabolic for daughter hypha<br><u>Apical extension</u>: constant during branching | <u>Location and dynamics</u>: parental Spitzenkörper progresses unchanged, the daughter hypha forms its own Spitzenkörper early and centrally | <u>Orientation</u>: parallel to the hyphal axes<br><u>Distribution</u>: axially symmetrical<br><u>Dynamics</u>: populations relatively constant |

**Table 1.**
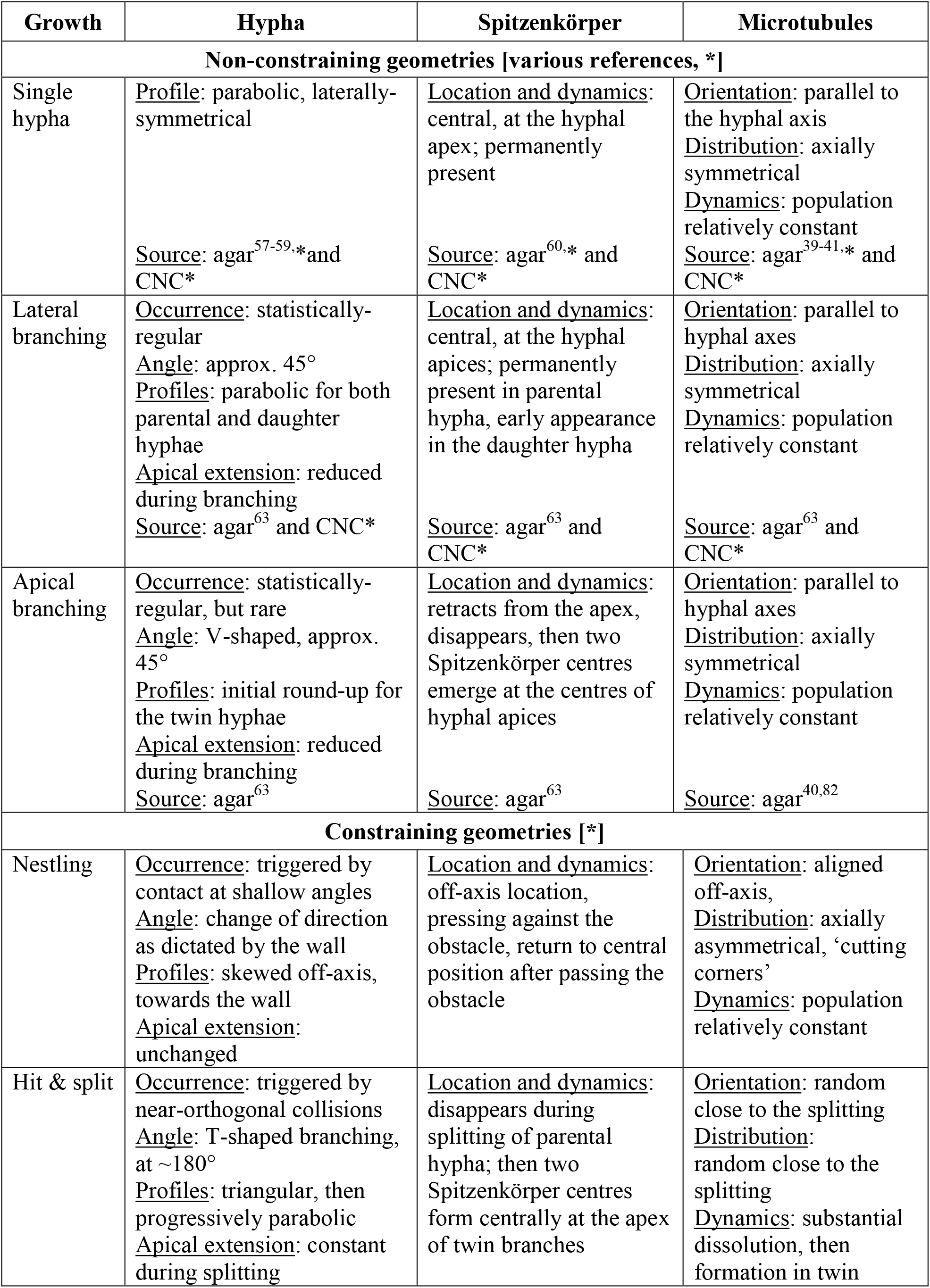
Intracellular processes in the apical region involved in the growth of *Neurospora crassa* in open and constraining environments ([*] indicates results from the present study)

| Growth | Hypha | Spitzenkörper | Microtubules |
| --- | --- | --- | --- |
| **Non-constraining geometries [various references, \*]** |  |  |  |
| Single hypha | <u>Profile</u>: parabolic, laterally-symmetrical<br><u>Source</u>: agar<sup>57-59,*</sup> and CNC* | <u>Location and dynamics</u>: central, at the hyphal apex; permanently present<br><u>Source</u>: agar<sup>60,*</sup> and CNC* | <u>Orientation</u>: parallel to the hyphal axis<br><u>Distribution</u>: axially symmetrical<br><u>Dynamics</u>: population relatively constant<br><u>Source</u>: agar<sup>39-41,*</sup> and CNC* |
| Lateral branching | <u>Occurrence</u>: statistically-regular<br><u>Angle</u>: approx. 45°<br><u>Profiles</u>: parabolic for both parental and daughter hyphae<br><u>Apical extension</u>: reduced during branching<br><u>Source</u>: agar<sup>63</sup> and CNC* | <u>Location and dynamics</u>: central, at the hyphal apices; permanently present in parental hypha, early appearance in the daughter hypha<br><u>Source</u>: agar<sup>63</sup> and CNC* | <u>Orientation</u>: parallel to hyphal axes<br><u>Distribution</u>: axially symmetrical<br><u>Dynamics</u>: population relatively constant<br><u>Source</u>: agar<sup>63</sup> and CNC* |
| Apical branching | <u>Occurrence</u>: statistically-regular, but rare<br><u>Angle</u>: V-shaped, approx. 45°<br><u>Profiles</u>: initial round-up for the twin hyphae<br><u>Apical extension</u>: reduced during branching<br><u>Source</u>: agar<sup>63</sup> | <u>Location and dynamics</u>: retracts from the apex, disappears, then two Spitzenkörper centres emerge at the centres of hyphal apices<br><u>Source</u>: agar<sup>63</sup> | <u>Orientation</u>: parallel to hyphal axes<br><u>Distribution</u>: axially symmetrical<br><u>Dynamics</u>: population relatively constant<br><u>Source</u>: agar<sup>40,82</sup> |
| **Constraining geometries [\*]** |  |  |  |
| Nestling | <u>Occurrence</u>: triggered by contact at shallow angles<br><u>Angle</u>: change of direction as dictated by the wall<br><u>Profiles</u>: skewed off-axis, towards the wall<br><u>Apical extension</u>: unchanged | <u>Location and dynamics</u>: off-axis location, pressing against the obstacle, return to central position after passing the obstacle | <u>Orientation</u>: aligned off-axis,<br><u>Distribution</u>: axially asymmetrical, ‘cutting corners’<br><u>Dynamics</u>: population relatively constant |
| Hit & split | <u>Occurrence</u>: triggered by near-orthogonal collisions<br><u>Angle</u>: T-shaped branching, at ~180°<br><u>Profiles</u>: triangular, then progressively parabolic<br><u>Apical extension</u>: constant during splitting | <u>Location and dynamics</u>: disappears during splitting of parental hypha; then two Spitzenkörper centres form centrally at the apex of twin branches | <u>Orientation</u>: random close to the splitting<br><u>Distribution</u>: random close to the splitting<br><u>Dynamics</u>: substantial dissolution, then formation in twin |

The present study shows that the intracellular processes involved in the growth of *Neurospora crassa* in constraining geometries are triggered by the presence of, and modulated by the type of obstacles encountered by hyphae. Of the two important behavioural traits of *Neurospora crassa* growth in constraining environments^26^, directional memory appears to be the result of the Spitzenkörper functioning as a compass preserving the initial direction of growth, and pressing against opposing obstacles encountered at a shallow angle of attack, then returning to the initial direction when the blocking obstacle is overtaken and the contact with the hypha ceases. This compass-like dynamic memory is further stabilised by the structuring of the microtubules in the wake of the trajectory of the Spitzenkörper, resulting in the characteristic ‘cutting corners’ feature of the microtubule cytoskeleton in meandering channels. Directional memory, evidenced as a behavioural trait in few fungal species,^25,26^ could provide biological benefits for the filamentous fungi growing and foraging in geometrically heterogeneous environments. Indeed, stochastic simulations showed that supressing directional memory in *Pycnoporus cinnabarinus*^25^ increases the probability of hyphae being trapped in a network. Furthermore, the *Neurospora crassa* ro-1 mutant, which did not display directional memory, presented a considerably more even distribution of the hyphal mass in networks, and consequently a considerably lower capacity for exiting from complex geometries than the wild-type *Neurospora crassa^26^*

In contrast with the intracellular processes involved in directional memory, the Spitzenkörper-microtubules system does not appear to determine the direction of obstacle-induced branching. Indeed, in the hit & split events, at the critical point of apical splitting, both the Spitzenkörper and the microtubules are absent. Furthermore, in lateral branching events triggered by the availability of lateral free space, during or after tight geometrical constrainment, and although the Spitzenkörper-microtubules system is present in the early stages of formation of the daughter branch, directional memory is unable to dictate its direction of growth. Arguably, the only driving force of the extension of the resulting hyphae, and thus of the obstacle-induced branching, is the isotropic turgor pressure. The presence of obstacle-induced branching in the same species exhibiting directional memory^25-27^ suggests that this behavioural trait also brings biological benefits. Indeed, stochastic simulations^25^ demonstrated that obstacle-induced branching leads to a higher capacity of evading complex networks, but with a lesser relative benefit than directional memory. Consequently, it appears that *Neurospora crassa* has evolved ‘hard wired’ intracellular processes responsible for directional memory and obstacle-induced branching, respectively, with the former being the main driver for the negotiation of complex networks, and the latter a fall-back mechanism when directional memory is turned-off during near-orthogonal collisions, or when it cannot operate due to the tight constraining in tight geometries.

Aside of the interest in the fundamentals of intracellular mechanisms involved in fungal growth, this study could have further impact in several directions, not exhaustively mentioned below.

- From a methodological perspective, purposefully-designed, optically transparent PDMS microfluidics structures, in conjunction with advanced microscopy imaging, can be used in fundamental microbiology studies by triggering, with temporal and spatial precision, biomolecular events which are modulated by the cellular interaction with the solid environment. This experimental methodology can be used for the further exploration of other elements controlling the fungal growth in confined spaces, in particular the role of actin structures, not covered in the present study.
- PDMS-made microfluidic devices can be designed to mimic fungal environments, with experiments revealing insight relevant to various environmental, industrial, and medical applications, including fungal pathogenicity in animals and plants. For instance, the PDMS mechanical strength can be adjusted to allow the estimation of the forces applied by fungi in various environmental conditions, via the measurement of resultant deformations, as already demonstrated for yeast.^68^ Alternatively, the design of the PDMS structures could mimic the structure of the walls of the plant or animal tissue to allow the study of fungal invasion.
- The confinement imposed on the growth of filamentous fungi can be used for various approaches of biologically-driven computation. For instance, it was shown^27^ that the genetically-engineered, cytoskeleton defective mutant of *Neurospora crassa*, which produces short branches preferentially at 90°, can solve orthogonal mazes better than the wild type *Neurospora crassa*, which is biologically ‘programmed’ to branch at 45°. Furthermore, as the space searching natural algorithms used by fungi have been demonstrated as being more efficient than some artificial ones^28^, it is possible to use fungi, either wild type, or better genetically-engineered, to solve very complex physical networks encoding combinatorial mathematical problems, as proposed before,^69^ and recently demonstrated.^70^ Alternatively, the nuclear dynamics in *Neurospora crassa*^71^ could be ‘streamlined’ in networks mimicking real complex transportation webs, thus allowing the exploration of traffic optimisation,^72,73^ a conceptual framework demonstrated for *Physarum polycephalum^74^*

### Conclusions

The study of the intracellular processes in somatic hyphae of *Neurospora crassa* that respond actively to geometrical constraints imposed by a PDMS-based microfluidic structure revealed how the Spitzenkörper-microtubules system is responsible for the directional memory in navigating confining networks when hyphae encounter obstacles at shallow angles of contact. This study also revealed that the Spitzenkörper-microtubules system is not modulating the obstacle-induced hyphae collide near-orthogonally with obstacles blocking their growth, suggesting that turgor pressure is the remaining candidate for the driving force. Finally, when free space becomes available laterally from tightly-constraining channels, Spitzenkörper-microtubules system-controlled directional memory cannot operate, also leaving turgor pressure as the last possible driving force for hyphal lateral branching. The present results can impact on further fundamental studies regarding the intracellular processes driving the fungal growth in confined environments, and on various environmental, industrial, and medical applications, as diverse as fungal pathogenicity in plants, animals and humans, to biologically-driven computation.

## Methods

### Microfabrication and experimental setup

The microfluidic network is illustrated in Figure 1 and Supplementary Figure SI 01. Its dimensions, i.e., height of 10 μm, and channel widths ranging from 2 to 100 μm were designed to present various level of constrainment to fungal growth, from tight-constraining in channels with widths smaller than the hyphal diameter, i.e., 5-7 μm, to confined, but non-constraining chambers, with dimensions of 100 x 100 x 10 μm. The artificial environments were fabricated using a two-component polymer, poly(dimethyldisiloxane) (PDMS, Sylgard 184, Dow Corning) using a well-established procedure.^26^ Benefits of using PDMS include low fabrication costs, non-toxicity, good biocompatibility, chemical inertness, and optical transparency for wavelengths as low as 280 nm.^75-79^ Briefly, the fabrication involved the moulding of a degassed PDMS mixture of the pre-polymer and curing agent (10:1, w/w) onto a microstructured silicon wafer, at 65°C for a duration in excess of 8 hours. After hydrophilisation via exposure to UV/ozone, the PDMS stamps were irreversibly fixed onto a microscope cover slip. Lateral openings in the structure allowed the introduction of the growth medium, fungal hyphae, and fluorescent dyes. Fungal inoculation was achieved by placing an agar plug, extracted from a zone with young hyphae, e.g., the peripheral growth zone of a colony, upside down next to a lateral channel opening. The device was then attached to a microscope slide marked with spacers for accurate positioning on a microscope stage. Hyphal confinement within channels ensured that the hyphae remained within the working distance of the microscope objective while enabling sufficient gas exchange over long periods of time, thus avoiding the need for perfusion with oxygenated nutrient broth, as required in agar.^60^,^63^

The microfluidic network design allowed the investigation of fungal behaviour in the following scenarios (Supplementary Figure SI 33, from top to bottom): (a) *virtually no mechanical confinement*, wherein hyphae with a diameter of 5-7 μm grow in the 10 μm gap between the glass coverslip and the PDMS ‘ceiling’), similar to agar; (b) *parallel 1D confinement*, wherein hyphae progress along a wall in the observation plane; (c) *2D confinement*, wherein hyphae grow while being constrained between two walls that are perpendicular to the observation plane; and (d) *orthogonal or angled 1D confinement*, wherein hyphae encounter a wall at near-normal incidence. In many instances, the hyphae encounter the wall at a shallower angle (e.g., 45° or less, relative to the surface), which results in a parallel 1D confinement. Additionally, in the case of 2D confinement, the channels can be given various widths and shapes (e.g., straight, zig-zagged, or bent at various angles).

### Fungal species, growth media, staining

The *Neurospora crassa rid (RIP4) mat a his-3+::Pccg-1-Bml+sgfp+* mutant strain (henceforth *“Neurospora crassa* GFP”; FGSC #9519) was obtained from the Fungal Genetics Stock Center (School of Biological Sciences, University of Missouri, Kansas City, MO, USA). The *Neurospora crassa GFP* mutant was constructed^41^ to express intrinsically GFP-labelled microtubules while maintaining a growth pattern similar to that of the wild type. The strain was cultured on 1% w/v malt extract agar (Merck), which was also used for medium filling the microfluidics structures. Prior to each experiment, the fungal strains were subcultured on fresh malt extract agar plates and incubated at room temperature (21°C ± 2°C).

The FM4-64 dye (Invitrogen Ltd. (Paisley, UK) was used as a marker for Spitzenkörper^80^. A 20-μl droplet of an 8 μM FM4-64 solution was applied onto a microscope coverslip before placing an agar slab, excised from the margin of the growing colony, upside-down onto the droplet. To avoid an overlay of the dynamics of the dye loading and of the Spitzenkörper, imaging was performed at least one hour after loading the hyphae with the dye.

### Time-lapse microscopy and image analysis

Live-cell imaging of hyphal growth was performed with an inverted laser-scanning microscope (Zeiss Axio Observer Z1 with LSM 5 Exciter RGB, Carl Zeiss, Göttingen, Germany) with photomultiplier detectors. Samples were excited with 488 nm and 543 nm lasers, and the emitted light was passed through a bandpass filter (505-530 nm) and a 650 nm long-pass filter. To reduce photobleaching and phototoxic effects, the laser intensity and laser scanning time were kept to a minimum (0.7 - 2.4 % laser energy, 0.75- to 23-second frame scans). Fluorescence and bright-field time-lapse images were captured simultaneously and analysed using image processing software (Zen 2008, Carl Zeiss, Göttingen, Germany). Fiji^81^ was used for image overlay and quantitative image analysis. RETRAC 2.10.0.5 (freeware from Dr. Nick Carter, University of Warwick, UK) was used for frame-by-frame tracking and calculating cytoskeletal and hyphal kinetics.

### Growth experiments on agar and microfluidics structures

Control measurements for fungal growth in non-constraining environments were performed on 1% w/v malt extract media using somatic hyphae at the edges of the colony. The leading hyphae, i.e., wide hyphae showing rapid cytoplasmic flow,^82^ rarely entered the microfluidic structures and were therefore omitted. For the somatic hyphae, ‘subapical compartments’ were characterised by an increased nuclear density approximately 60 μm from the extreme apex. Hyphal growth rates were established by tracking the position of the extreme hyphal apices in subsequent frames. To measure the cytoskeletal alignment within hyphae, tangents were fitted manually to microtubules, and the respective local hyphal longitudinal (i.e., polarisation) axes and intersection angles were measured. To measure the rates of microtubule polymerisation within the apical compartment, the positions of individual filament ends were tracked frame-by-frame.

The parameters of the obstacle-induced apical hit & split included the time elapsed from the impact to the establishment of the daughter hyphae and the maximum size of the formed bulges immediately before the re-establishment of polarised growth. The hyphal diameter was measured at the time of collision with the obstacle. The maximum bulge size was measured by overlaying the frame of collision with the frame in which the growth pattern of the daughter bulges changed to polarised extension and determining the difference in the apical cell wall location on both sides of the hypha.

### Statistical analysis

Statistica 7.1 (Statsoft Inc., OK, USA) and GraphPad Prism 6.01 (GraphPad Software Inc., CA, USA) were used for statistical analysis and correlation tests. Statistical analyses included calculating the mean and standard deviation values of parameters measured, i.e., position, alignment with the hyphal axis, polymerisation rate for microtubules, times before reappearance of the Spitzenkörper, and hyphal bulge dimensions, over the total number *n* data points. Statistical analyses included all accumulated data from at least 20 separate experiments (unless otherwise stated). GraphPad prism was used to perform a Mann-Whitney test comparing the apical and subapical distributions of the microtubule polymerisation rates and the microtubule alignments to the polarisation axis respectively.

## Acknowledgements

Financially supported by the European Union Seventh Framework Programme (FP7/2007-2011) under Grant Agreements 228971 [Molecular Nano Devices (MONAD)]; by a Research Project Grant from Leverhulme Trust; and by grants from the Defense Advanced Research Projects Agency under Grant Agreements N66001-03-1-8913, and HR0011-16-2-0028. We thank Dr. Adam Hendricks, from McGill University, for insightful suggestions, and Drs. Abraham P. Lee and Lisen Wang, from University of California, Irvine, for the fabrication of the microfluidics masters.

## Author contributions

MH conceived and conducted the experiments, and performed the image acquisition and analysis. OK contributed to the image and statistical analyses. CE contributed to the analysis of the biological data. DVN conceived the experiments, contributed to the image analysis and coordinated the project. MH and DVN wrote the paper.

